# Paradoxical activation of SREBP1c and *de novo* lipogenesis by hepatocyte-selective ACLY depletion in obese mice

**DOI:** 10.1101/2022.03.21.485183

**Authors:** Batuhan Yenilmez, Mark Kelly, Guofang Zhang, Nicole Wetoska, Olga R. Ilkayeva, Kyounghee Min, Leslie Rowland, Chloe DiMarzio, Wentao He, Naideline Raymond, Lawrence Lifshitz, Meixia Pan, Xianlin Han, Jun Xie, Randall H. Friedline, Jason K. Kim, Guangping Gao, Mark A. Herman, Christopher B. Newgard, Michael P. Czech

## Abstract

Hepatic steatosis associated with high fat diets, obesity and type 2 diabetes is thought to be the major driver of severe liver inflammation, fibrosis, and cirrhosis. Cytosolic acetyl-coenzyme A (AcCoA), a central metabolite and substrate for de novo lipogenesis (DNL), is produced from citrate by ATP-citrate lyase (ACLY) and from acetate through AcCoA synthase short chain family member 2 (ACSS2). However, the relative contributions of these two enzymes to hepatic AcCoA pools and DNL rates in response to high fat feeding is unknown. We report here that hepatocyte-selective depletion of either ACSS2 or ACLY caused similar 50% decreases in liver AcCoA levels in obese mice, showing that both pathways contribute to generation of this DNL substrate. Unexpectedly however, the hepatocyte ACLY depletion in obese mice paradoxically increased total DNL flux measured by D2O incorporation into palmitate, while in contrast ACSS2 depletion had no effect. The increase in liver DNL upon ACLY depletion was associated with increased expression of nuclear sterol regulatory element-binding protein 1c (SREBP1c) and of its target DNL enzymes. This upregulated DNL enzyme expression explains the increased rate of palmitate synthesis in ACLY depleted livers. Furthermore, this increased flux through DNL may also contribute to the observed depletion of AcCoA levels due to its increased conversion to Malonyl CoA (MalCoA) and palmitate. Together, these data indicate that in HFD fed obese mice, hepatic DNL is not limited by its immediate substrates AcCoA or MalCoA, but rather by activities of DNL enzymes.

## INTRODUCTION

One of the debilitating co-morbidities of type 2 diabetes in obesity is nonalcoholic steatohepatitis (NASH), characterized by severe liver inflammation and fibrosis that can lead to cirrhosis and the need for liver transplantation (1–8). NASH is driven by several cell types within the liver, including pro-inflammatory cells and stellate cells that secrete collagens and other extracellular matrix proteins that lead to fibrosis(3–12). Development of hepatic steatosis often precedes NASH, associated with increased rates of fatty acid synthesis (*de novo* lipogenesis, DNL) and lipid sequestration within lipid droplets(13–17). Based on the identification of polymorphisms in genes such as patatin-like phospholipase domanin containing protein 3 (PNPLA3), known to regulate lipid metabolism but also associated with progression of NASH, it is thought that hepatic steatosis is an important therapeutic target pathway for alleviating NASH(18–21). Enzyme targets for limiting hepatic steatosis have been proposed and investigated, including diacylglycerol acyl transferase 2 (DGAT2) (22–29), which catalyzes the final step of triglyceride synthesis, and AcCoA carboxylase (ACC1) (30, 31), the enzyme that converts acetyl CoA (AcCoA) to malonyl CoA (MalCoA) in the DNL pathway. Pharmacologic suppression of these enzymes has elicited decreases in hepatic steatosis as well as some indicators of NASH in humans (24, 32). Taken together, these considerations highlight the importance of gaining a more complete understanding of the dynamics of hepatic lipid synthesis in response to high fat/high sucrose diet (HFD) feeding, as well as in overt obesity and type 2 diabetes.

A central metabolic intermediate in lipid metabolism is AcCoA, which in the cytosol is the substrate for both fatty acid and cholesterol synthesis, and in mitochondria serves as an intermediate of glucose and fatty acid oxidation (33). Cytosolic AcCoA has two well studied sources—citrate, which yields AcCoA and oxaloacetate when cleaved by ACLY, and acetate, which is converted to AcCoA by ACSS2. Recent studies have demonstrated that fructose supplementation via the drinking water induces hepatic DNL through multiple mechanisms including induction of hepatic lipogenic enzymes, and by metabolism of fructose by the microbiome to generate acetate, which then serves as the substrate for DNL via its conversion to AcCoA by ACSS2 (34–36) The importance of microbial acetate as a source of fructose-derived lipogenic substrate is supported by experiments showing that knockout of ACLY in mice has no impact on total rates of hepatic DNL in response to fructose feeding (34). The relative contributions of ACLY and ACSS2 to AcCoA production and DNL flux has not been studied in the context of the most common experimental model of obesity, its induction by feeding with HFD. In humans, inhibition of ACLY by the small molecule drug bempedoic acid decreases blood lipids without decreasing liver fat, but combination therapies that simultaneously target ACLY and ACSS2 have not been reported (37–39).

Based on these considerations, we have performed a study of the effects of liver-specific suppression of ACLY (ACLY LKO) or ACSS2 (ACSS2 LKD) expression alone, or of both genes combined (Double), on hepatic DNL and levels of key metabolic intermediates including AcCoA, MalCoA and acetate in mice fed HFD. Several surprising findings emerged: 1. While hepatocyte AcCoA levels were decreased by depletion of either ACLY or ACSS2 alone, or in combination, MalCoA levels were unchanged in response to any of these maneuvers, indicating that neither ACLY nor ACSS2 are required for maintaining levels of this immediate DNL precursor in HFD-fed obese mice; 2. There is a compensatory increase in nuclear SREBP1c and expression of enzymes in the DNL pathway such as fatty acid synthase (FASN) and stearoyl-Coenzyme A desaturase 1 (SCD1) when hepatic ACLY is depleted, leading to a paradoxical increase in DNL in the absence of ACLY in HFD-fed obese mice, and 3. The upregulation of DNL enzymes occurring in response to hepatic ACLY depletion is associated with a compensatory increase in ACSS2 and a corresponding decrease in circulating acetate levels.

Altogether, these studies reveal unexpected features of hepatic DNL flux in obese mice and provide a framework for understanding mechanisms that link AcCoA producing enzymes to control of more distal enzymes in the DNL pathway.

## RESULTS

In order to determine the relative contributions of metabolic flux through ACLY versus ACSS2 to form AcCoA and fuel DNL in hepatocytes *in vivo,* we used mice floxed flanking exon 17-19 of the *acly* gene and injected with either AAV engineered for hepatocyte selective Cre expression (pAAV-TBG-PI-Cre) to delete liver ACLY, or AAV engineered for hepatocyte selective expression of an artificial micro-RNA directed against ACSS2 (pAAV-TBG-amiRACSS2) to achieve liver ACSS2 depletion (Fig. 1A). To obtain combined depletion of ACLY and ACSS2, both AAV constructs were injected simultaneously. To establish the utility of these vectors, *acly*-floxed mice fed on CHOW diet were injected with these AAV constructs or a control AAV and sacrificed 9 weeks later for analysis (Fig. 1B). Immunoblotting analysis of livers from these mice revealed the expected depletion of ACLY, ACSS2 or both proteins dependent on the vectors administered (Fig. 1C). In addition, immunoblots of brown, epididymal and inguinal adipose tissue showed no decreases in ACLY or ACSS2 expression, confirming the liver specificity of the gene silencing provided by the AAV constructs (Fig.S1). No difference in body weight was noted in response to treatment with any of the AAV vectors (data not shown). Hepatic AcCoA levels did not change significantly in response to suppression of either ACLY or ACSS2 alone, but were reduced ~50% in response to combined depletion of hepatic ACLY plus ACSS2 (Fig. 1D). These data validate the efficacy of the AAV constructs to cause significant depletion of the targeted enzymes, and demonstrate that with CHOW feeding, the mice are able to maintain normal hepatic AcCoA levels by alternate routes when either ACLY or ACSS2 levels are diminished, but not when both are depleted.

**Figure 1:**
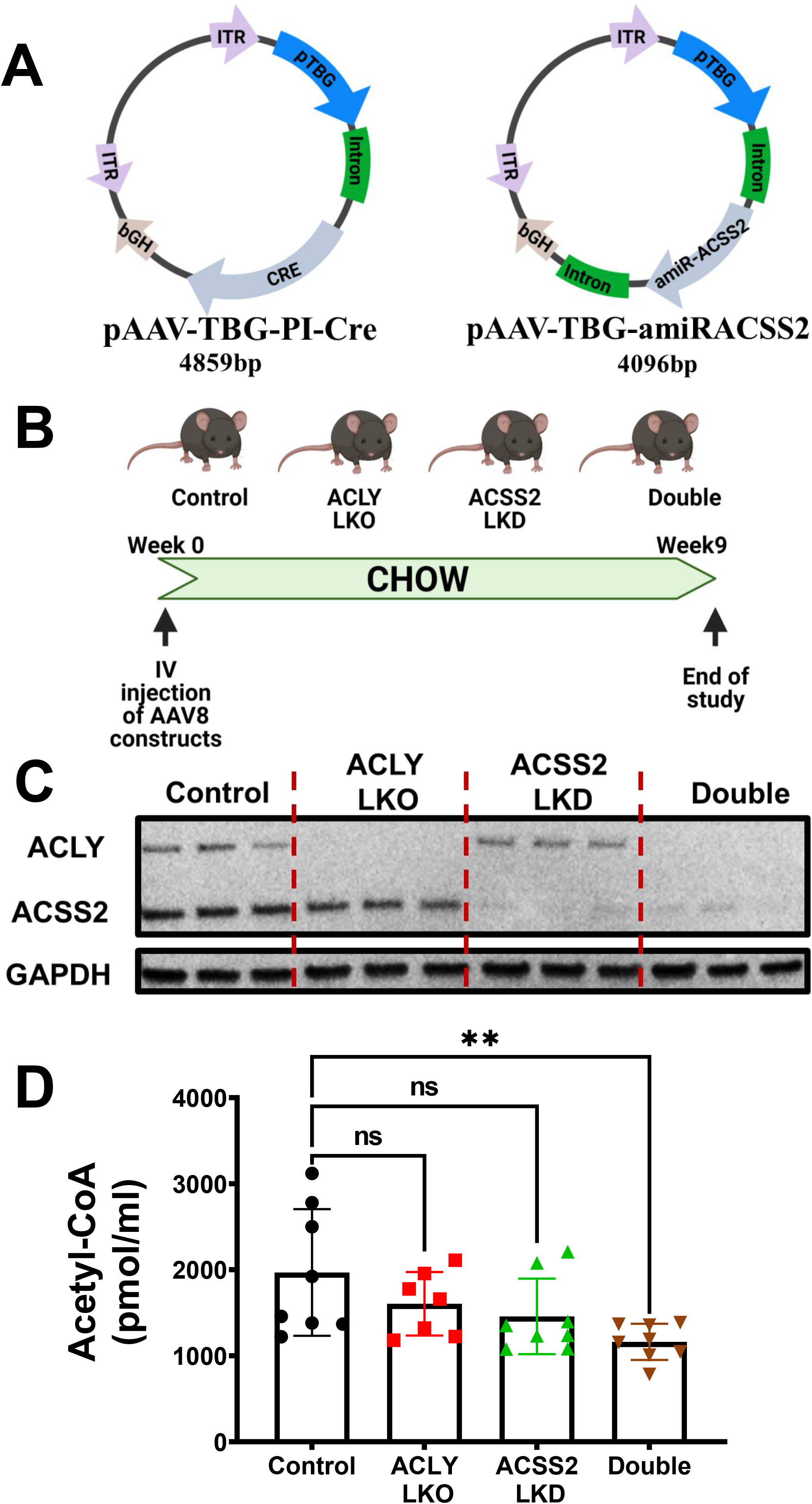
Depletion of both ACLY and ACSS2 in hepatocytes of CHOW fed mice decreases the hepatic AcCoA pool. Eight week old male *Acly* fl/fl C57BL6/J mice (n=10) fed on CHOW were injected IV with AAV8-TBG-PI-Cre to deplete ACLY, AAV8-TBG-amiRACSS2 to deplete ACSS2, or with both vectors to deplete both enzymes in a hepatocyte-specific manner. Nine weeks after injection mice were sacrificed and tissues harvested. **(A)** Schematic rendering of viral vectors used to deplete ACLY (expressing CRE under TBG promoter) and ACSS2 (expressing artificial miRNA targeting *Acss2* mRNA under TBG promoter) **(B)** Study plan for investigation of hepatocyte specific depletion of ACLY and/or ACSS2 in *Acly* fl/fl C57BL6 mice on chow diet **(C)** Confirmation of lowering of ACLY and ACSS2 protein levels by immunoblotting. **(D)** Hepatic AcCoA levels measured by mass spectrometry (ns: Not significant, *: p<0.05, **:p<0.005, ***:p<0.0005, ****:p<0.00005)

Figure 2A depicts the experimental protocol used to address the key questions of our study concerning regulation of DNL in HFD-fed obese mice. Following injection of the AAV constructs, mice were fed CHOW for one week, and then switched to a diet containing 60% fat (HFD) for 8 additional weeks prior to sacrifice. No changes in body weight (Fig. 2B) or food intake (Fig. 2C) were observed in mice injected with the ACLY or ACCS2 AAV vectors relative to mice injected with control AAV. Treatment of HFD-fed mice with pAAV-TBG-PI-Cre caused near complete suppression of ACLY mRNA (Fig. 2D) and protein (Fig 2E,F) levels, both when administered alone or in conjunction with the pAAV-TBG-amiRACSS2 vector that depletes ACSS2. However, in contrast to what was observed in CHOW fed mice (Fig.1), depletion of hepatocyte ACLY caused significant upregulation of ACCS2 expression. Moreover, while injection of the pAAV-TBG-amiRACSS2 vector alone caused a strong depletion of *Acss2* mRNA and protein in the obese mice, this suppression was less effective when combined with ACLY depletion, likely due to the compensatory upregulation phenomenon (Fig. 2D,E&F). The levels of ACSS2 protein in double knockout mice were similar to those in mice treated with the control AAV vector but were well below levels observed in the ACLY LKO mice.

**Figure 2:**
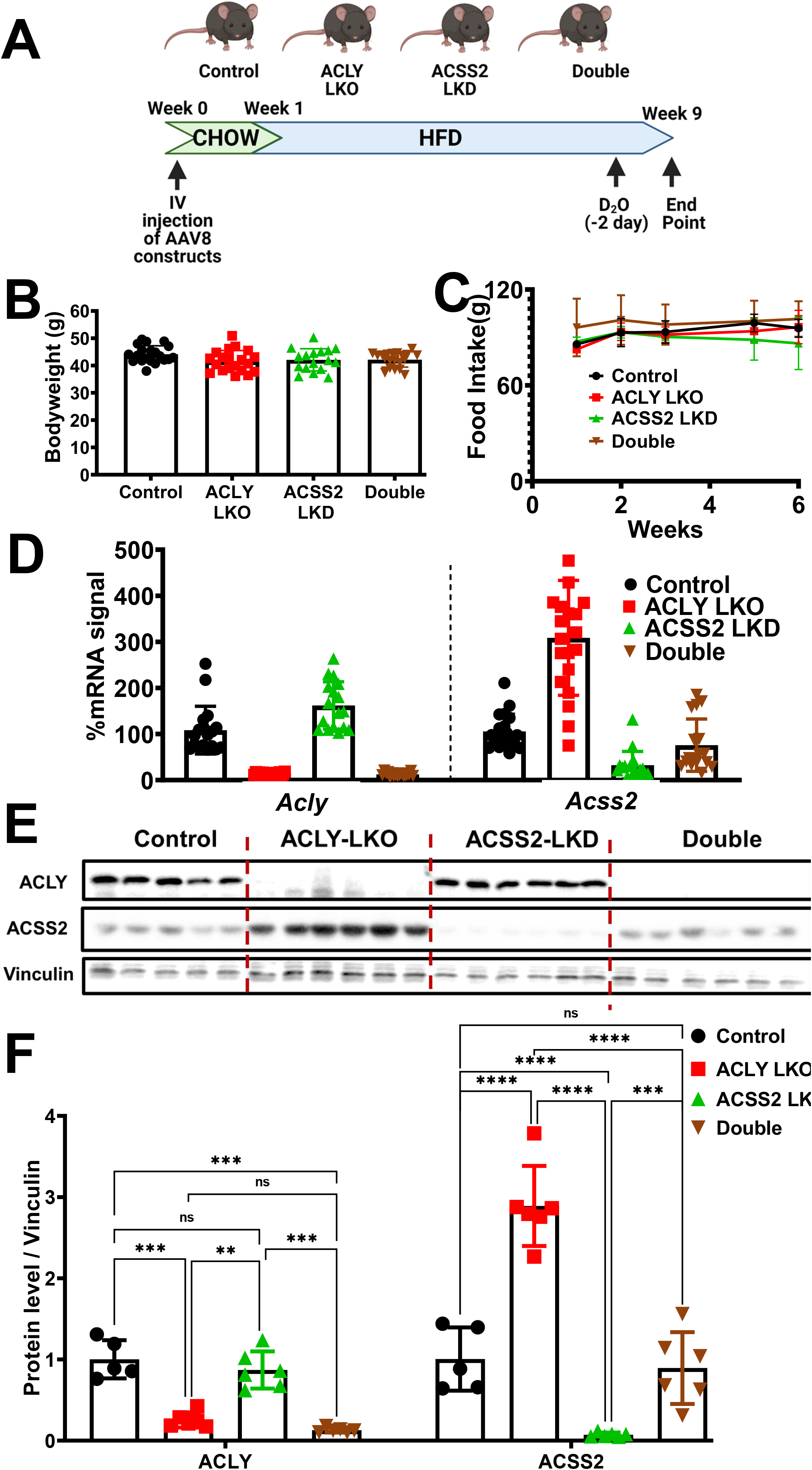
Hepatic depletion of ACLY causes a compensatory upregulation of ACSS2 expression in mice on HFD. Eight week old male *Acly* fl/fl C57BL6/J mice (n=20) on CHOW were injected with viral particles via IV. A week after injection, mice were switched to 60% HFD. After 8 weeks of HFD feeding mice were sacrificed and tissues harvested. Two days before the study end-date mice were injected with D_2_O (25ul/g dose) and their drinking water was switched to 6% D_2_O to allow DNL measurements. **(A)** Summary of study plan **(B)** Bodyweight measurements at the end of 8 weeks of HFD feeding **(C)** Food intake measurements for the duration of HFD feeding. Meaurements of ACLY and ACSS2 **(D)** mRNA and **(E)** protein levels in liver **(F)** Quantification of ACLY and/or ACSS2 protein levels. (ns: Not significant, *: p<0.05, **:p<0.005, ***:p<0.0005, ****:p<0.00005)

Next, metabolite levels and metabolic flux through DNL were assessed in two separate cohorts of HFD-fed, obese mice, and the results of the two studies were combined to provide the data in Fig. 3. Under HFD conditions, knockdown of either ACLY or ACSS2 in hepatocytes led to a significant decrease in AcCoA levels, with a trend for further decline in mice with combined knockdown of both enzymes (Fig. 3A). Notably, the compensatory increase in ACSS2 expression in the ACLY LKO condition did not restore AcCoA levels in the absence of ACLY. That ACSS2 is active under these conditions is verified by analysis of plasma acetate (Fig 3B), which was inversely proportional to ACSS2 expression. Notably, plasma acetate levels were also decreased in response to combined suppression of ACLY and ACSS2, suggesting that even though ACSS2 protein was not elevated in this condition compared to mice treated with the control AAV vector, flux through this enzyme and its consumption of acetate may have increased. Remarkably, the fall in AcCoA levels elicited by knockdown of ACLY, ACSS2 or both enzymes was not accompanied by a decrease in MalCoA levels (Fig 3C). Also unanticipated, newly synthesized palmitate and total palmitate levels were increased in response to ACLY LKO, either alone or when combined with ACSS2 suppression, whereas ACSS2 LKD caused a modest decrease in these outcomes, despite the decrease in AcCoA levels in response to all of these maneuvers (Figs. 3B,D,E).

**Figure 3:**
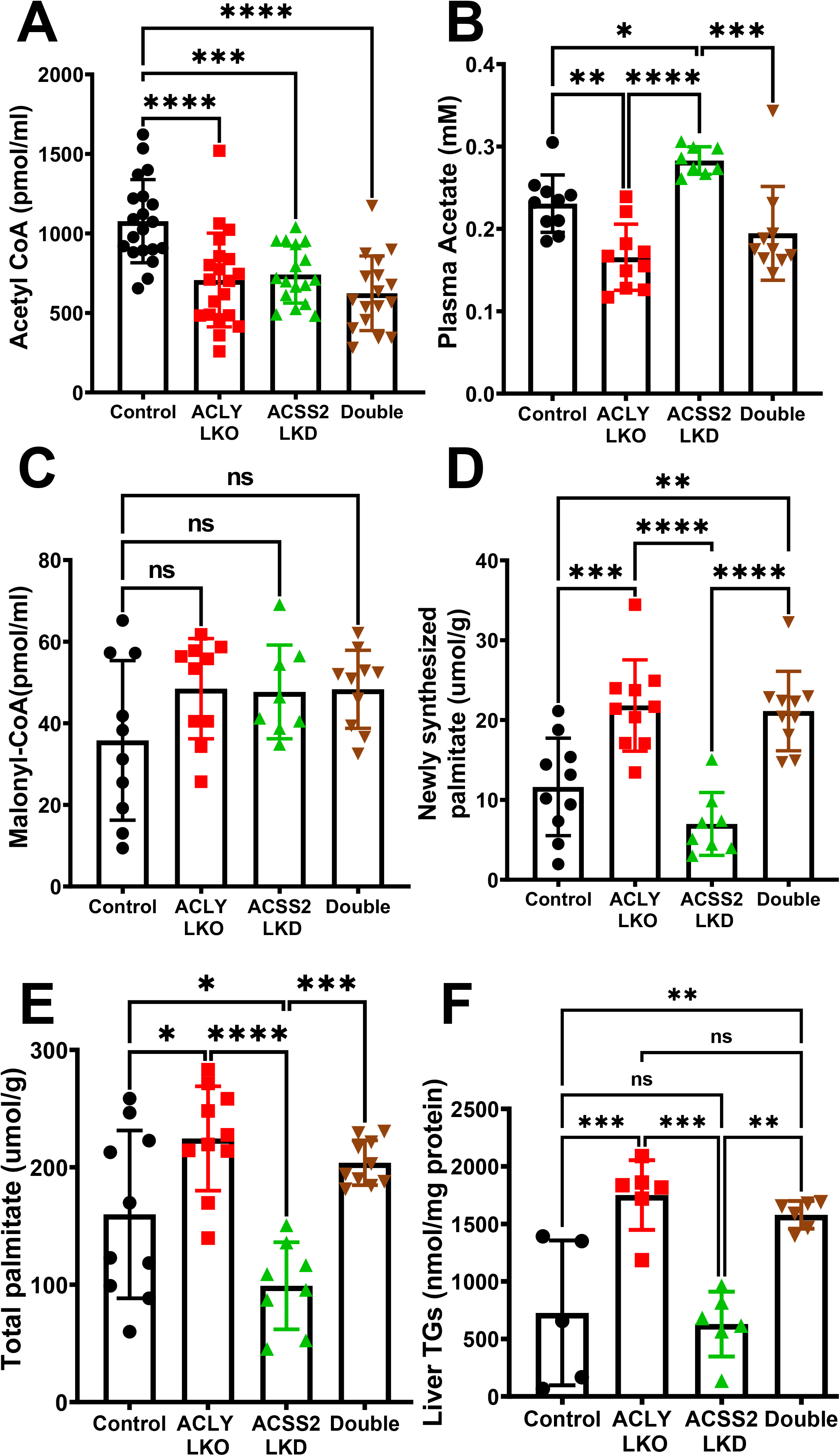
Hepatic depletion of ACLY alone or both ACLY and ACSS2 decrease hepatic AcCoA levels but increase DNL and liver TG in mice fed HFD. Liver homogenates were processed to measure specific metabolite changes resulting from depletion of ACLY and/or ACSS2. **(A)** Hepatic AcCoA levels **(B)** Plasma Acetate levels **(C)** Hepatic MalCoA levels measured by LC-MS/MS **(D)** Hepatic newly synthesized palmitate levels (measured by deuterium incorporation from D_2_O into liver palmitate)**(E)** Hepatic total palmitate levels **(F)** Liver TG levels per mg of protein. (ns: Not significant, *: p<0.05, **:p<0.005, ***:p<0.0005, ****:p<0.00005)

Total liver triglyceride (TG) levels also followed the newly synthesized palmitate and total palmitate trend. Hepatic TG levels were significantly increased by ACLY depletion or double depletion compared to the ACSS2 LKD and control group (Fig 3F). None of these changes in hepatic lipid biosynthesis were reflected in changes in plasma lipids (Fig.S2). These data indicate that AcCoA levels do not determine MalCoA levels or rates of DNL under HFD fed conditions in mice. Instead, the metabolite that most strongly correlated, in an inverse fashion, with DNL was plasma acetate, which was reduced in response to ACLY depletion (alone or in concert with ACSS2 knockdown), and slightly increased in response to ACSS2 LKD alone.

We next explored the mechanisms underlying the paradoxical increase in DNL engendered by ACLY LKO, occurring in the face of lowered AcCoA levels and unchanged MalCoA levels. To this end, we measured expression of the genes encoding key enzymes in the DNL pathway, as well as transcription factors known to control their expression (Fig. 4). At the mRNA level, two enzymes in the DNL pathway were found to be significantly elevated in ACLY depleted livers—*Fasn and Scd1* (Fig 4A). In contrast, ACSS2 depletion alone, which had no significant effect on newly synthesized palmitate (Fig 3D), also had no effect on *Fasn* or *Scd1* mRNA levels (Fig 4A). Immunoblot analyses confirmed the increase in FASN and SCD1 at the protein level, and also revealed an increase in ACC1, both total amount and it’s phosphorylated, inactivated form in the ACLY LKO condition, resulting in no net change in the inactive to active ACC1 ratio. (Fig 4B, C). Interestingly, depletion of ACSS2 in addition to ACLY LKO abrogated the increase in DNL enzyme mRNA levels but not protein levels. These data suggest that hepatic DNL in HFD fed obese mice is not limited by or dependent on AcCoA and MalCoA levels, but rather is regulated by altered expression and activities of DNL enzymes such as FASN and SCD1.

**Figure 4:**
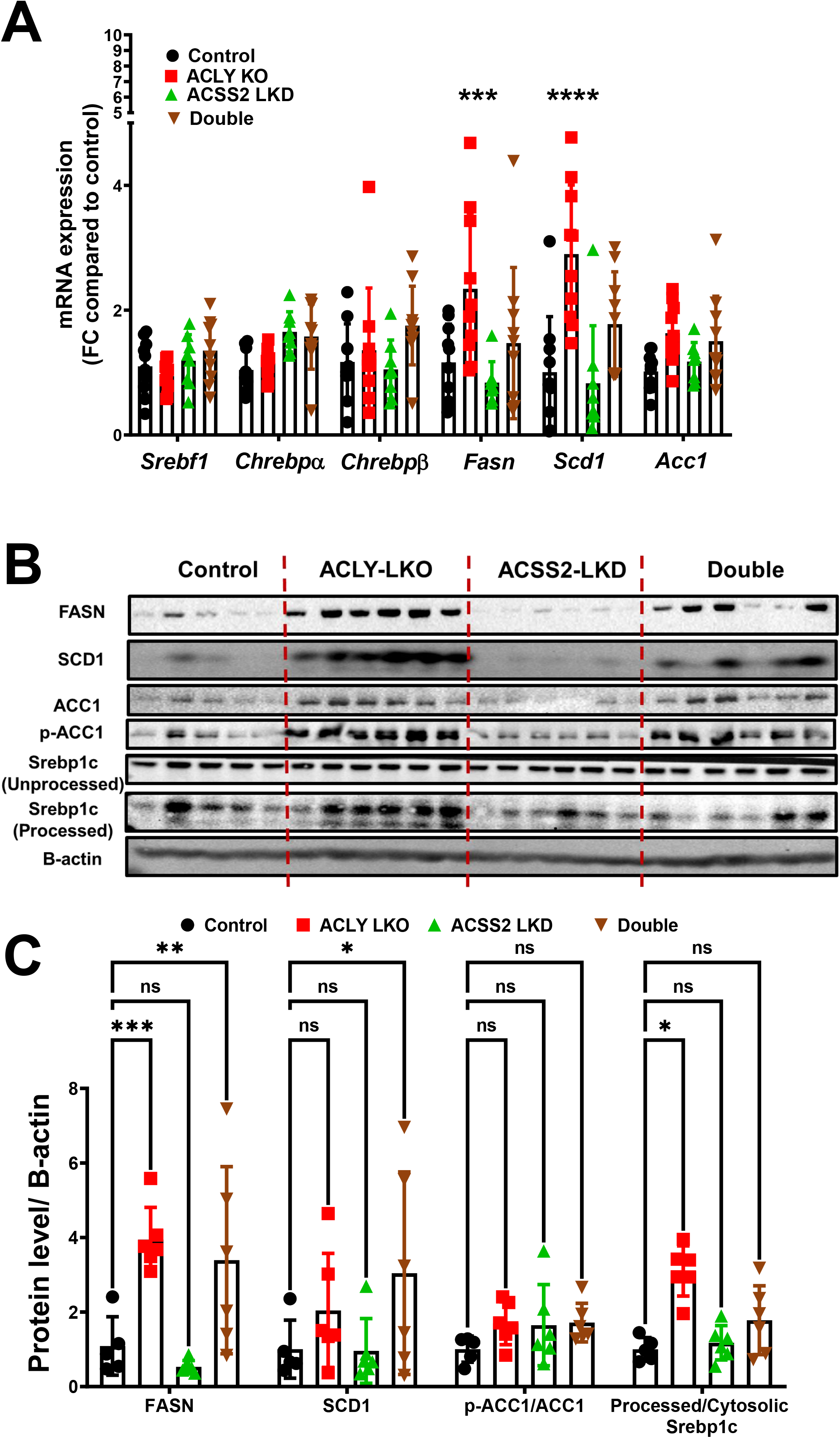
Hepatic depletion of ACLY but not of ACSS2 results in upregulation of enzymes in the DNL pathway processed (active) Srebp1c protein levels. Livers from the HFD fed mouse groups were processed for investigation of **(A)** mRNA and **(B)** protein expression changes between the groups. **(C)** Quantification of protein levels between groups. (ns: Not significant, *: p<0.05, **:p<0.005, ***:p<0.0005, ****:p<0.00005)

Since it is known that the transcription factors ChREBP as well as SREBP1c regulate the expression of DNL enzymes in liver, we analyzed expression of these transcription factors in our genetically engineered HFD-fed mice. While no difference in expression levels of these transcription factors at the mRNA level were observed, we found a clear effect on levels of the nuclear-localized form of SREBP1c. Figures 4B and 4C show that hepatic ACLY LKO but not ACSS2 LKD causes increased processing of SREBP1c to the nuclear form, an effect not observed when ACLY LKO was combined with ACSS2 LKD. These data suggest that ACLY depletion causes upregulation of SREBP1c processing to its activated form, associated with upregulation of ACSS2 as well as SCD1 and FASN. Although this effect is not significant when ACLY is depleted in combination with ACSS2 depletion, the trend is still evident and the DNL enzymes are upregulated in this condition (Fig. 4C).

We applied correlation analyses of data from individual mice to investigate how Srebp1 processing and DNL enzyme expression may be related to the ACLY and ACSS2 perturbations (Fig. 5). This analysis confirmed the lack of correlation between AcCoA levels and newly synthesized palmitate, but showed a significant correlation between plasma acetate levels and DNL (Fig 5A,B). Moreover, there were strong correlations between FASN protein expression (Fig 5C) and SREBP1c processing (Fig 5C) with DNL flux. Importantly, plasma acetate concentrations correlated inversely with FASN expression (Fig 5D) and with the nuclear form of SREBP1c (Fig 5D), consistent with the concept that acetate levels are an indicator of SREBP1c activity, hepatic DNL enzyme expression, and hepatic DNL flux.

**Figure 5:**
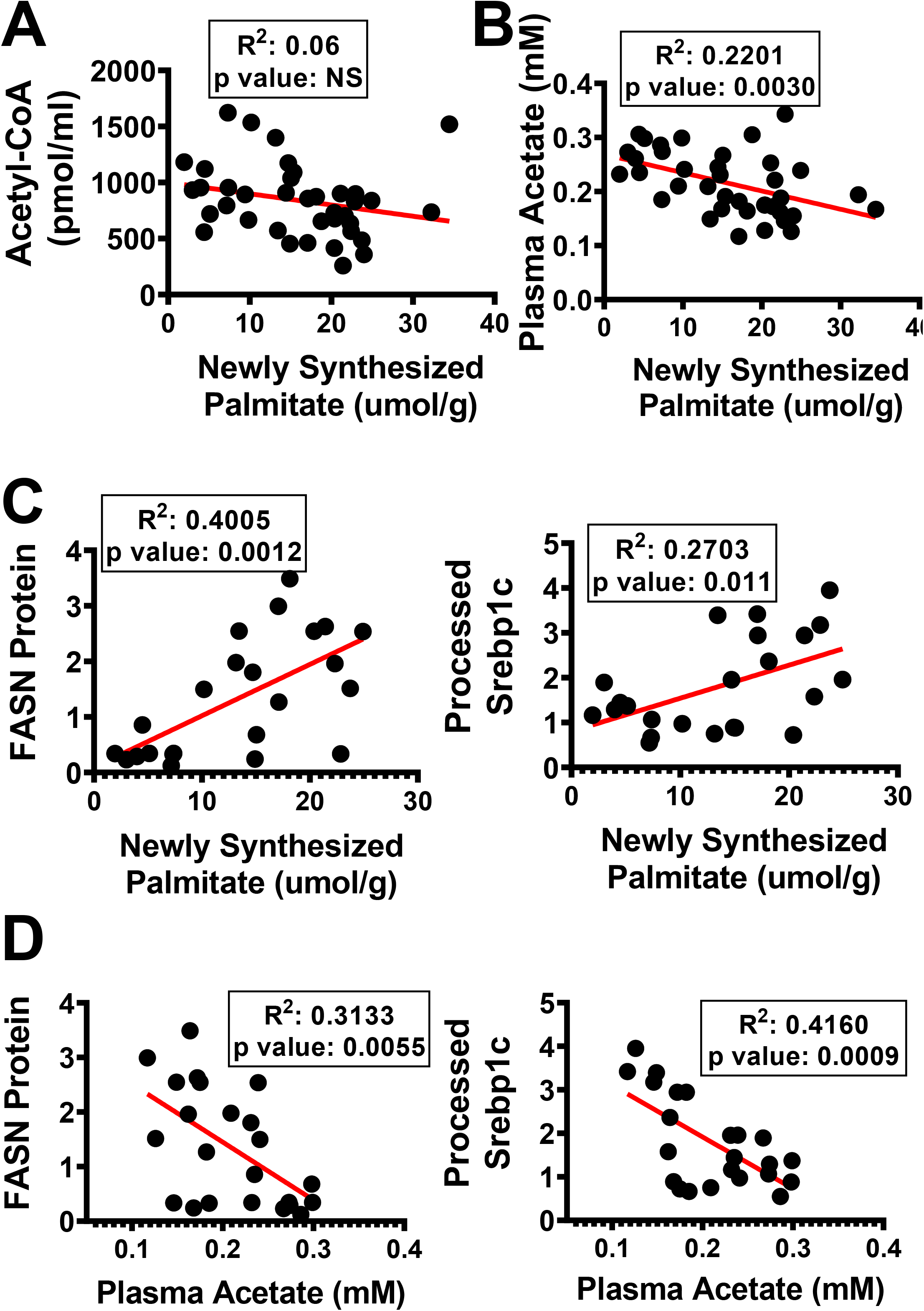
Newly synthesized palmitate levels (DNL) and plasma acetate correlate with FASN expression levels but not AcCoA. Correlation graphs of **(A)** AcCoA versus newly synthesized palmitate, **(B)** Plasma acetate versus newly synthesized palmitate **(C)** FASN and processed Srebp1c protein levels versus newly synthesized palmitate **(D)** FASN and processed Srebp1c protein levels versus plasma acetate. (ns: Not significant, *: p<0.05, **:p<0.005, ***:p<0.0005, ****:p<0.00005)

In addition, lipidomics analysis showed significantly increased diglyceride accumulation as well as fatty acyl chains in TGs in ACLY LKO and double depletion groups compared to ACSS2 LKD and control groups (Fig. S2A&B). Interestingly the increase in newly synthesized palmitate, liver TGs and diglyceride accumulation were inversely correlated with phospholipid biosynthesis, suggesting that DNL is shunted towards TG biosynthesis compared to phospholipid biosynthesis under conditions where ACLY is absent (Fig. S3C,D&E).

## DISCUSSION

A primary question addressed in this study is the degree to which the two AcCoA synthesis pathways catalyzed by ACLY vs ACSS2 contribute to steady state levels of AcCoA in livers of HFD-fed, obese mice. The importance of this question is reinforced by several perspectives: 1. AcCoA is a major substrate in the pathways of DNL and cholesterol synthesis, both of which are up-regulated in obesity to contribute to metabolic disease, 2. AcCoA is a substrate for histone and other protein acetylation reactions that control gene expression and enzyme function, and 3. Effects of single and double depletions of hepatocyte ACLY and ACSS2 on levels of key DNL intermediates such as AcCoA, MalCoA and acetate, and flux through the DNL pathway, have not been investigated in the most commonly used animal model of obesity, the HFD-fed mouse. To perform these experiments in the most rigorous manner, AAV vectors were employed expressing TBG promoter-based constructs to assure hepatocyte-selective depletion of ACLY and ACSS2 (Fig. 1). The results indicate that ACLY and ACSS2 both contribute significantly to AcCoA levels in hepatocytes of HFD-fed, obese mice, as evidenced by significant decreases in content following depletion of each enzyme alone or both in combination (Figs. 2,3 & S1). In contrast, in mice on CHOW diet hepatic AcCoA levels were unchanged in response to suppression of either ACLY or ACSS2 alone and declined only when both ACLY and ACSS2 were suppressed (Fig.1D). These data show that HFD fed, obese mice, unlike lean mice, require both ACLY and ACSS2 to maintain their normal hepatic AcCoA levels.

It is important to note that there are some limitations in the interpretations that we can make in these studies. Even with the double depletion of ACLY and ACSS2 in hepatocytes, no more than a 50% decrement in hepatic AcCoA was observed in either the CHOW or HFD fed mice (Figs 1 and 3). In obese mice, the partial decrease in AcCoA might be explained by the lack of complete elimination of ACSS2 in the double depletion condition (Fig. 2E), although this does not explain the modest decrease in AcCoA levels in CHOW-fed mice with complete suppression of both hepatic enzymes (Fig 1C). There are several possible reasons for this modest decrease in the measured AcCoA levels when both enzymes are missing. First, other pathways of cytosolic AcCoA generation may be at play. For example, it has been proposed that cytosolic AcCoA can be regenerated from acetylcarnitine by a putative cytosolic carnitine acetyltransferase, which has been reported to be present in mammalian tissue, albeit not yet in liver (40, 41). Second, the extracts assayed here are from whole liver, and therefore include other cell types that also contain AcCoA such as endothelial cells and macrophages that are not depleted of ACLY or ACSS2, given that we specifically targeted hepatocytes through TBG promoter-driven AAV constructs. Third, cytosolic AcCoA derived from ACLY and ACSS2 represents only one compartment of AcCoA in the hepatocyte, whereas significant AcCoA pools in other cellular compartments such as mitochondria and peroxisomes are well established (41–47). In mitochondria, AcCoA is produced by glucose, fatty acid, and amino acid oxidation (42, 43, 46–52). In peroxisomes, AcCoA is also generated as a result of active fatty acid oxidation by a pathway defined by the rate limiting enzyme ACOX1(53). Finally, cytosolic AcCoA may be derived via export from the nucleus(46, 54, 55), although this pool is expected to also require ACLY and ACSS2 as does the cytosolic pool. Taken together, these considerations indicate that the roughly 50% decrease in AcCoA observed in the whole livers of HFD fed mice depleted of ACLY or ACSS2 represent minimal estimates of the reduction in the cytosolic AcCoA pool in hepatocytes.

In the case of single ACLY LKO, a decrease of about 40% in AcCoA occurs despite a strong compensatory upregulation of ACSS2 protein to approximately 3 times its normal level (Fig. 2D&E). Yet, when this ACSS2 compensation is blocked in the double knockdown, i.e., when ACLY is depleted and ACSS2 remains at basal levels (Fig 2D&E), AcCoA levels are lowered to a similar extent as in the ACLY LKO alone. While this result seems to suggest that ACSS2 may not contribute much to the AcCoA levels, ACSS2 LKD alone causes a 40% decrease in AcCoA, affirming it’s significant contribution to the pool (Fig 3A). Reconciliation of these findings may be reached by our observation that in HFD-fed, obese mice, ACLY LKO causes a large increase in expression of DNL enzymes (Fig 4) and in DNL flux to increase palmitate synthesis (Fig 3D&E). This increased DNL flux secondary to the upregulation of downstream DNL enzymes likely exerts a “pull” on the AcCoA pool (Fig 3A). This occurs in addition to the presumed decreased generation of AcCoA from ACLY depletion.

Interestingly, MalCoA levels remain unchanged in the face of ACLY or ACSS2 depletion, even though utilization of this intermediate for DNL was clearly increased in ACLY LKO mice. Taken together, our results suggest that ACSS2 is the predominant contributor to AcCoA generation in HFD-fed, obese mice since ACSS2 depletion reduces AcCoA levels to an extent equal to the ACLY LKO condition, but with no change in DNL flux (Figs. 3 and 4). This concept is consistent with work indicating a major role for acetate in liver lipogenesis as recently published by others using different mouse models (34, 56, 57). Furthermore, it is consistent with the marked decrease in plasma acetate that occurs in conjunction with the upregulation of ACSS2 in the ACLY LKO condition, indicating significant flux of acetate into DNL (Fig 3B). A strong correlation between the decrease in plasma acetate and increase in DNL activity (Fig. 5B) is also noted, further supporting the role of acetate as a major substrate for DNL in our HFD-fed mouse model of obesity.

The results described above raise an important mechanistic question: How does hepatocyte ACLY depletion in HFD-fed, obese mice result in increased expression of DNL enzymes? We initially found no changes in mRNA expression levels of two of the major transcription factors that regulate enzymes of lipid biosynthesis, SREBP1c (58–60) and ChREBP (61–64). However, further investigation revealed a clear increase in levels of the processed, nuclear-localized and active form of SREBP1c protein in livers of obese ACLY LKO mice (Fig.4B). The increase in mature SREBP1c levels strongly correlated with DNL rate (Fig 5C). Thus, our results suggest that ACLY LKO in hepatocytes of obese mice causes proteolytic processing of SREBP1c to its transcriptionally active form through an unknown mechanism. Two pathways known to modulate SREBP1c are mTORC1 (65–67) and AMPK (56, 68). However, we did not observe a change in mTORC1 activity in response to ACLY LKO, as measured by levels of p70S6K or p4-EBP. We also were unable to detect significant changes in levels of p-AMPK levels in the ACLY LKO condition (data not shown). Thus, the mechanism connecting ACLY expression and regulation of SREBP1c processing will require further investigation.

It is noteworthy that studies in other mouse model systems have not revealed a connection between ACLY LKO and increased DNL activity that we report here in the HFD-obese mouse model. Wang et al. (69) used adenovirus-shRNA to deplete ACLY in db/db mice (a genetic model of obesity) but fed the mice CHOW rather than HFD diet. In that setting, depletion of ACLY caused a reduction in total liver AcCoA, similar to our findings in HFD-obese mice (Fig 3), but contrary to our findings, they also reported reduced DNL as measured by D2O incorporation into newly synthesize palmitate, reduced liver TGs, and reduced expression of DNL enzymes in the ACLY knockdown condition (69). On the other hand, Zhao et al. (34) used mice on a high fructose diet to study ACLY and ACSS2 depletion in liver and reported that fructose-mediated lipogenesis is driven by acetate produced in the gut (34). When labeled glucose was used to measure DNL rates in that study, depletion of ACLY abolished DNL, as expected since glucose carbons must pass through ACLY to promote DNL. However, when D2O was used for measurements of DNL, no effect of ACLY depletion was observed, in contrast with the marked increase we report here (34). It is possible that in addition to the differences in dietary model (high fructose vs high fat), differences in the time course of the experiment and other study details such as their use of albumin promoter-driven Cre recombinase to elicit ACLY depletion, and the extent of compensatory ACSS2 induction in their model, may help to explain the contrasting results.

Although data in humans at the level of detail described in our studies is not available, the ACLY inhibitor bempedoic acid has been used clinically in human subjects (38, 70–74). In phase 3 clinical trials, bempedoic acid reduced the mean LDL cholesterol level by up to 30% (74) in patients with hypercholesterolemia and by around 18% in patients on statin medications (73). Moreover, the safety and efficacy of bempedoic acid were consistently favorable following one year of administration (73), and FDA approval was obtained in February 2020. In our plasma analyses of total cholesterol (Fig S3A), no differences were observed under conditions of single or double depletions of ACLY and ACSS2 compared to control groups (Fig.S2). One possible reason for difference is that bempedoic acid treatment has been shown to induce upregulation of LDL receptor (72), which in turn would increase the uptake of lipoproteins from the circulation. Other possible explanations for the divergent results include the species difference (mouse versus human) and our hepatocyte-specific deletion of ACLY in mice versus orally delivered bempedoic acid in human subjects that may be more widely distributed to other tissues. Bempedoic acid is converted to its bioactive form by very long chain acyl-CoA synthase (ACSVL1), which is purported to be liver-specific (38, 70–74). However, single cell RNA sequencing and the human protein atlas data base reveals considerable ACSVL1 mRNA and protein presence in other metabolically active tissues such as kidney and gastrointestinal tissues (75–77). Potential systemic effects as a consequence of ACLY inhibition in kidney and the gastrointestinal track remain to be investigated. Moreover, there is no data showing reduction of liver DNL in humans in response to bempedoic acid administration, and therefore it is not yet possible to compare our results in mice to human subjects.

In summary, we demonstrate that in HFD-fed, obese mice both ACLY and ACSS2 contribute to the AcCoA pool in hepatocytes, although the ACSS2 pathway may predominate. Importantly, depletion of hepatic ACLY in these mice triggers a mechanism whereby SREBP1c is activated, and DNL-related enzymes are upregulated to increase the DNL rate. Thus, distal DNL enzymes rather than AcCoA-generating enzyme are the primary regulators of hepatic palmitate synthesis in obese mice during HFD feeding.

## MATERIALS AND METHODS

### Animal Studies

Animal experiments were performed in accordance with animal care ethics approval and guidelines of University of Massachusetts Medical School Institutional Animal Care and Use Committee (IACUC, protocol number A-1600-19). For in vivo studies *Acly* floxed mice were purchased from Taconic Biosciences. Eight-week-old, male, *Acly* floxed mice were injected with corresponding AAV8 constructs via intravenous injection to achieve hepatocyte-specific depletion of ACLY and/or ACSS2. One week after the administration of the AAV8 constructs, these mice either stayed on standard CHOW or were switched to HFD (Research Diets D12492) for 8 weeks. Two days prior to the end point of the study, mice were pre-bled and then injected with 25ul/g of bodyweight D2O along with supplementation of 6% D2O into the drinking water. Blood samples were taken at 1 day prior to the end point as well as terminally for measurement of D2O enrichment in the circulation.

### Generation of hepatocyte specific AAV8 construct

The artificial miRNA against mouse Acss-2 was designed as previous described (78) and the synthesized gBlock was incorporated after the liver specific TBG promoter in pAAV-TBG-PI vector plasmid (79) by Gibson assembly. rAAV8 was produced by transient HEK 293 cell transfection and CsCl sedimentation by the University of Massachusetts Medical School Viral Vector Core, as previously described (80). Vector preparations were determined by ddPCR, and purity was assessed by 4%–12% SDS-acrylamide gel electrophoresis and silver staining (Invitrogen)

### RNA isolation and RT-qPCR

Frozen liver tissue punches (25-50 mg) were homogenized in trizol using the Qiagen TissueLyser II. Chloroform was added to the homogenate and centrifuged for 15 min at max speed. The clear upper layer was added to an equal volume of 100% isopropanol and incubated for 1 hour at 4° C. After 10 min of centrifugation at max speed, the supernatant was discarded and 70-75% ethanol was added to wash the pellet. After 15 min centrifugation at max speed, the supernatant was discarded, and the pellet was briefly dried in the hood before being resuspended in ddH2O. RNA concentration was then measured by the Thermo Scientific NanoDrop2000 spectrophotometer. cDNA was synthesized from 1 μg of total RNA using iScript cDNA Synthesis Kit (BioRad) and BioRad T100 thermocycler. Quantitative RT-PCR was performed using iQ SybrGreen Supermix on the BioRad CFX96 C1000 Touch Thermal Cycler and analyzed as previously described(81).

### Immunoblotting

For protein expression analyses, frozen liver tissue (~25 mg) was homogenized by the Qiagen TissueLyser II in a sucrose buffer (250 mM sucrose, 50 mM tris-Cl pH 7.4) with 1:100 protease inhibitor (Sigma-Aldrich). The tissue lysates were denatured by boiling, separated on a 4-15% sodium dodecyl sulfate/ polyacrylamide gel electrophoresis gel (BioRad) and transferred to a nitrocellulose membrane (BioRad). The membrane was blocked with Tris-buffered saline with Tween (TBST) containing 5% milk for 1 hour at room temperature and incubated with primary antibodies; ACLY, ACSS2, FASN, SCD1, ACC1, p-ACC1, GAPDH, β-actin purchased from Cell Signaling, or SREBP1c antibody purchased from Millipore. The blot was washed in TBST for an hour, incubated at room temperature with corresponding second antibody at room temperature for 30 min, washed again, and incubated with ECL (Perkin Elmer) and visualized with the ChemiDox XRS+ image-forming system.

### Lipidomics Analysis

Lipid species in liver samples were analyzed using multidimensional mass spectrometry-based shotgun lipidomic analysis (82). In brief, each liver tissue sample homogenate containing 0.5 mg of protein (determined with a Pierce BCA assay) was accurately transferred to a disposable glass culture test tube. A pre-mixture of lipid internal standards (IS) was added prior to conducting lipid extraction for quantification of the targeted lipid species. Lipid extraction was performed using a modified Bligh and Dyer procedure, and each lipid extract was reconstituted in chloroform:methanol (1:1, v:v) at a volume of 400 μL/mg protein (83).

For shotgun lipidomics, the lipid extract was further diluted to a final concentration of ~500 fmol total lipids per μL. Mass spectrometric analysis was performed on a triple quadrupole mass spectrometer (TSQ Altis, Thermo Fisher Scientific, San Jose, CA) and a Q Exactive mass spectrometer (Thermo Scientific, San Jose, CA), both of which were equipped with an automated nanospray device (TriVersa NanoMate, Advion Bioscience Ltd., Ithaca, NY) as described (84). Identification and quantification of lipid species were performed using an automated software program (85). Data processing (e.g., ion peak selection, baseline correction, data transfer, peak intensity comparison and quantitation) was performed as described(85). The results were normalized to the protein content (nmol lipid/mg protein).

### Metabolomics

Liver acyl CoA esters were analyzed as previously described (86, 87) by flow injection analysis using positive electrospray ionization on Xevo TQ-S, triple quadrupole mass spectrometer (Waters, Milford, MA) employing methanol/water (80/20, v/v) containing 30 mM ammonium hydroxide as the mobile phase. Spectra were acquired in the multichannel acquisition mode monitoring the neutral loss of 507 amu. Heptadecanoyl CoA was employed as an internal standard. The endogenous CoAs were quantified using calibrators prepared by spiking liver homogenates with authentic CoAs (Sigma, St. Louis, MO) having saturated acyl chain lengths C2-C18. Corrections for the heavy isotope effects, mainly ^13^C, to the adjacent m+2 spectral peaks in a particular chain length cluster were made empirically by referring to the observed spectra for the analytical standards.

Malonyl CoA was extracted with 0.3 M perchloric acid and analyzed by LC-MS/MS using a previously published method (88). The extracts were spiked with ^13^C2-Acetyl-CoA (Sigma, MO, USA), centrifuged, and filtered through Millipore Ultrafree-MC 0.1 μm centrifugal filters before being injected onto a Chromolith FastGradient RP-18e HPLC column, 50 x 2 mm (EMD Millipore) and analyzed on a Waters Xevo TQ-S triple quadrupole mass spectrometer coupled to a Waters Acquity UPLC system (Waters, Milford, MA).

### *De novo* lipogenesis measurements

*De novo* lipogenesis was measured as previously described (49). Total palmitic acid labeling assay in the liver was assayed by GC-MS. Briefly, 20 mg liver tissue was homogenized in 1 ml KOH/EtOH (EtOH 75%) and incubated at 85 °C for 3 hours, and 200 μl of 1 mM [^13^C16]palmitate was added to samples as internal standard after cool down. Extracted palmitate acid was mixed with 50 μl N-tert-Butyldimethylsilyl-N-methyltrifluoroacetamide (TBDMS) at 70 °C for 30 minutes, and the TBDMS-derivatized samples were analyzed with an Agilent 5973N-MSD equipped with an Agilent 6890 GC system, and a DB-17MS capillary column (30 m x 0.25 mm x 0.25 um). The mass spectrometer was operated in the electron impact mode (EI; 70 eV). The temperature program was as follows: 100°C initial, increase by 15°C/min to 295°C and hold for 8 min. The sample was injected at a split ratio of 10:1 with a helium flow of 1 ml/min. Palmitate-TBDMS derivative eluted at 9.7 min, and the m/z at 313, 314, and 319 were extracted for M0, M1, and M16 palmitate quantification. Stable isotope labeling was corrected for natural isotope abundance(89). Newly synthesized palmitic acid was calculated as: %newly synthesized palmitic acid labeling = total palmitic acid labeling /(plasma ^2^H_2_O labeling × 22)×100.

### Plasma acetate measurements

Acetate in plasma was quantified by LC-MS/MS method as described (90, 91), with modifications. 30 μl plasma was mixed with 30 μl of 200 μM [^2^H_5_]acetate as internal standard. Acetonitrile (1 ml) was added and the sample was centrifuged for 20 minutes at 8000×g to remove protein. The supernatant was transferred to a new Eppendorf vial and was dried completely under N_2_ gas. The dried residue was derivatized with 20 μl 120 mM 1-ethyl-3-(3-dimethylaminopropyl)carbodiimide hydrochloride, 20 μl 200 mM 3-nitrophenylhydrazine hydrochloride, and 50 μl LC-MS grade water at 40 °C for 30 minutes. The sampe was subjected to LC-MS/MS analysis, with ionization and fragmentation of acetate/[^2^H_5_]acetate optimized in negative ESI by QTRAP 6500^+^-MS (Sciex, Framinham, MA). A gradient LC method was developed with an Agilent Pursuit XRs 5 C18 column (150 ×2.0 mm, 5 μm). Mobile phase A was 98% water (LC-MS-grade) and 2% acetonitrile (LC-MS-grade) containing 0.1% formic acid. Mobile phase B was 98% acetonitrile and 2% water containing 0.1% formic acid. MS/MS ion transitions for acetate and [^2^H_5_]acetate were m/z 194/151 and 199/155, respectively. Data was analyzed by Analyst 1.6 software.

### Software & Statistics

All statistical analyses were performed using the GraphPad Prism 8 (GraphPad Software, Inc.). The data are presented as mean ± SEM. For analysis of the statistical significance between four or more groups, two-way ANOVA and multiple comparison t tests were used. Ns is nonsignificant (p > 0.05), *:p < 0.05, **:p < 0.005, and ***:p < 0.0005.

## Data Availability

The data that support the findings of this study are openly available upon request.

## Acknowledgements

We wish to thank members of our respective laboratories for helpful discussions, and Kerri Miller for excellent assistance throughout the project. This work was supported by National Institutes of Health grants DK103047 (to M.P.C.), DK121710 (to C.B.N and M.A.H.) and DK124723 (to C.B.N.). We also gratefully acknowledge generous funding through the Isadore and Fannie Foxman Chair in Medical Science (to M.P.C.), and pre-doctoral fellowship support to Batuhan Yenilmez by the American Heart Association (AHA Award #19PRE34460013).

## Author Contributions

B.Y., M.P.C and C.B.N designed the study, analyzed data and wrote the manuscript. B.Y., M.K, O.R.I, N.W, G.Z, K.M, C.D and N.D. performed most of the experiments and analyzed the data. O.R.I performed the metabolomics experiments. G.Z and W.Y. performed the DNL measurements and plasma acetate measurements. L.R. and M.H contributed to discussions with expert opinion on metabolism. L.L contributed to discussions on bioinformatic analyses. M.P. and X.H. performed the lipidomics analyses. J.X and G.G contributed with AAV generation and their expertise in AAV biology. Plasma cholesterol, FFAs and TG measurements were performed by the NIH funded Mouse Metabolic Phenotyping Center (NIH grant 5U2C-DK093000) at the University of Massachusetts Chan Medical School.

## REFERENCES

1. Younossi ZM, Koenig AB, Abdelatif D, Fazel Y, Henry L, and Wymer M. Hepatology. John Wiley and Sons Inc.; 2016:73–84.

2. Younossi ZM, Blissett D, Blissett R, Henry L, Stepanova M, Younossi Y, et al. Hepatology. John Wiley and Sons Inc.; 2016:1577–86.

3. Singh S, Khera R, Allen AM, Murad MH, and Loomba R. Hepatology. John Wiley and Sons Inc.; 2015:1417–32.

4. Musso G, Cassader M, Rosina F, and Gambino R. Diabetologia. Diabetologia; 2012:885–904.

5. Lomonaco R, Ortiz-Lopez C, Orsak B, Webb A, Hardies J, Darland C, et al. Hepatology. Hepatology; 2012:1389–97.

6. Friedman SL, Neuschwander-Tetri BA, Rinella M, and Sanyal AJ. Nature Medicine. Nature Publishing Group; 2018:908–22.

7. Estes C, Razavi H, Loomba R, Younossi Z, and Sanyal AJ. Hepatology. John Wiley and Sons Inc.; 2018:123–33.

8. Alexander M, Loomis AK, Van Der Lei J, Duarte-Salles T, Prieto-Alhambra D, Ansell D, et al. BMC Medicine. BioMed Central Ltd.; 2019.

9. Mahady SE, Webster AC, Walker S, Sanyal A, and George J. Journal of Hepatology. J Hepatol; 2011:1383–90.

10. Esler WP, and Bence KK. Cellular and Molecular Gastroenterology and Hepatology. Elsevier; 2019:247–67.

11. Drenth JPH, and Schattenberg JM. Expert Opinion on Investigational Drugs. Taylor and Francis Ltd.; 2020:1365–75.

12. Bence KK, and Birnbaum MJ. Molecular Metabolism. Elsevier GmbH; 2021:101143.

13. W J, and A S. Molecular metabolism. Mol Metab; 2020.

14. Schuster S, Cabrera D, Arrese M, and Feldstein AE. Nature Reviews Gastroenterology & Hepatology 2018 15:6. Nature Publishing Group; 2018:349–64.

15. Muthiah MD, and Sanyal AJ. Liver International. Blackwell Publishing Ltd; 2020:89–95.

16. M N, and A J S. Current hepatology reports. Curr Hepatol Rep; 2018:350–60.

17. Lambert JE, Ramos-Roman MA, Browning JD, and Parks EJ. Gastroenterology. W.B. Saunders; 2014:726–35.

18. Yang C, McDonald JG, Patel A, Zhang Y, Umetani M, Xu F, et al. Journal of Biological Chemistry. 2006:27816–26.

19. Wang Y, Kory N, BasuRay S, Cohen JC, and Hobbs HH. Hepatology. John Wiley & Sons, Ltd; 2019:2427–41.

20. Chamoun Z, Vacca F, Parton RG, and Gruenberg J. Biology of the Cell. John Wiley & Sons, Ltd; 2013:219–33.

21. BasuRay S, Wang Y, Smagris E, Cohen JC, and Hobbs HH. Proceedings of the National Academy of Sciences. National Academy of Sciences; 2019:9521–6.

22. Amin NB, Carvajal-Gonzalez S, Purkal J, Zhu T, Crowley C, Perez S, et al. Targeting diacylglycerol acyltransferase 2 for the treatment of nonalcoholic steatohepatitis. Science Translational Medicine. 2019;11(520):eaav9701.

23. Choi CS, Savage DB, Kulkarni A, Yu XX, Liu Z-X, Morino K, et al. Suppression of Diacylglycerol Acyltransferase-2 (DGAT2), but Not DGAT1, with Antisense Oligonucleotides Reverses Diet-induced Hepatic Steatosis and Insulin Resistance. Journal of Biological Chemistry. 2007;282(31):22678–88.

24. Loomba R, Morgan E, Watts L, Xia S, Hannan LA, Geary RS, et al. Novel antisense inhibition of diacylglycerol-acyltransferase 2 for treatment of non-alcoholic fatty liver disease: a multicentre, double-blind, randomised, placebo-controlled phase 2 trial. The Lancet Gastroenterology & Hepatology. 2020;5(9):829–38.

25. Yamaguchi K, Yang L, McCall S, Huang J, Yu XX, Pandey SK, et al. Inhibiting triglyceride synthesis improves hepatic steatosis but exacerbates liver damage and fibrosis in obese mice with nonalcoholic steatohepatitis. Hepatology. 2007;45(6):1366–74.

26. Yen C-LE, Stone SJ, Koliwad S, Harris C, and Farese RV, Jr. Thematic Review Series: Glycerolipids. DGAT enzymes and triacylglycerol biosynthesis. Journal of Lipid Research. 2008;49(11):2283–301.

27. Yenilmez B, Wetoska N, Kelly M, Echeverria D, Min K, Lifshitz L, et al. An RNAi therapeutic targeting hepatic DGAT2 in a genetically obese mouse model of nonalcoholic steatohepatitis. Mol Ther. 2021.

28. Yu XX, Murray SF, Pandey SK, Booten SL, Bao D, Song X-Z, et al. Antisense oligonucleotide reduction of DGAT2 expression improves hepatic steatosis and hyperlipidemia in obese mice. Hepatology. 2005;42(2):362–71.

29. Zammit Victor A. Hepatic triacylglycerol synthesis and secretion: DGAT2 as the link between glycaemia and triglyceridaemia. Biochemical Journal. 2013;451(1):1–12.

30. Goedeke L, Bates J, Vatner DF, Perry RJ, Wang T, Ramirez R, et al. Acetyl-CoA Carboxylase Inhibition Reverses NAFLD and Hepatic Insulin Resistance but Promotes Hypertriglyceridemia in Rodents. Hepatology. 2018;68(6):2197–211.

31. Kim C-W, Addy C, Kusunoki J, Anderson NN, Deja S, Fu X, et al. Acetyl CoA Carboxylase Inhibition Reduces Hepatic Steatosis but Elevates Plasma Triglycerides in Mice and Humans: A Bedside to Bench Investigation. Cell Metabolism. 2017;26(2):394–406.e6.

32. Calle RA, Amin NB, Carvajal-Gonzalez S, Ross TT, Bergman A, Aggarwal S, et al. ACC inhibitor alone or coadministered with a DGAT2 inhibitor in patients with non-alcoholic fatty liver disease: two parallel, placebo-controlled, randomized phase 2a trials. Nature Medicine. 2021;27(10):1836–48.

33. Choudhary C, Weinert BT, Nishida Y, Verdin E, and Mann M. The growing landscape of lysine acetylation links metabolism and cell signalling. Nat Rev Mol Cell Biol. 2014;15(8):536–50.

34. Zhao S, Jang C, Liu J, Uehara K, Gilbert M, Izzo L, et al. Dietary fructose feeds hepatic lipogenesis via microbiota-derived acetate. Nature. 2020;579(7800):586–91.

35. Perry RJ, Peng L, Barry NA, Cline GW, Zhang D, Cardone RL, et al. Acetate mediates a microbiome-brain-β-cell axis to promote metabolic syndrome. Nature. 2016;534(7606):213–7.

36. Herman MA, and Birnbaum MJ. Molecular aspects of fructose metabolism and metabolic disease. Cell Metab. 2021;33(12):2329–54.

37. Feng X, Zhang L, Xu S, and Shen A-z. ATP-citrate lyase (ACLY) in lipid metabolism and atherosclerosis: An updated review. Progress in Lipid Research. 2020;77:101006.

38. Ference BA, Ray KK, Catapano AL, Ference TB, Burgess S, Neff DR, et al. Mendelian Randomization Study of ACLY and Cardiovascular Disease. New England Journal of Medicine. 2019;380(11):1033–42.

39. Wei J, Leit S, Kuai J, Therrien E, Rafi S, Harwood HJ, et al. An allosteric mechanism for potent inhibition of human ATP-citrate lyase. Nature. 2019;568(7753):566–70.

40. Muoio DM, Noland RC, Kovalik JP, Seiler SE, Davies MN, DeBalsi KL, et al. Muscle-specific deletion of carnitine acetyltransferase compromises glucose tolerance and metabolic flexibility. Cell Metab. 2012;15(5):764–77.

41. Madiraju P, Pande SV, Prentki M, and Madiraju SR. Mitochondrial acetylcarnitine provides acetyl groups for nuclear histone acetylation. Epigenetics. 2009;4(6):399–403.

42. Patel MS, Nemeria NS, Furey W, and Jordan F. The pyruvate dehydrogenase complexes: structure-based function and regulation. J Biol Chem. 2014;289(24):16615–23.

43. Strittmatter L, Li Y, Nakatsuka NJ, Calvo SE, Grabarek Z, and Mootha VK. CLYBL is a polymorphic human enzyme with malate synthase and β-methylmalate synthase activity. Hum Mol Genet. 2014;23(9):2313–23.

44. Kunze M, Pracharoenwattana I, Smith SM, and Hartig A. A central role for the peroxisomal membrane in glyoxylate cycle function. Biochim Biophys Acta. 2006;1763(12):1441–52.

45. Chen Y, Siewers V, and Nielsen J. Profiling of cytosolic and peroxisomal acetyl-CoA metabolism in Saccharomyces cerevisiae. PLoS One. 2012;7(8):e42475.

46. Shi L, and Tu BP. Acetyl-CoA and the regulation of metabolism: mechanisms and consequences. Curr Opin Cell Biol. 2015;33:125–31.

47. Pietrocola F, Galluzzi L, Bravo-San Pedro José M, Madeo F, and Kroemer G. Acetyl Coenzyme A: A Central Metabolite and Second Messenger. Cell Metabolism. 2015;21(6):805–21.

48. McGarrah RW, Zhang G-F, Christopher BA, Deleye Y, Walejko JM, Page S, et al. Dietary branched-chain amino acid restriction alters fuel selection and reduces triglyceride stores in hearts of Zucker fatty rats. American Journal of Physiology-Endocrinology and Metabolism. 2020;318(2):E216–E23.

49. White PJ, McGarrah RW, Grimsrud PA, Tso SC, Yang WH, Haldeman JM, et al. The BCKDH Kinase and Phosphatase Integrate BCAA and Lipid Metabolism via Regulation of ATP-Citrate Lyase. Cell Metab. 2018;27(6):1281–93.e7.

50. Harris RA, Joshi M, Jeoung NH, and Obayashi M. Overview of the molecular and biochemical basis of branched-chain amino acid catabolism. J Nutr. 2005;135(6 Suppl):1527s–30s.

51. Violante S, Ijlst L, Ruiter J, Koster J, van Lenthe H, Duran M, et al. Substrate specificity of human carnitine acetyltransferase: Implications for fatty acid and branched-chain amino acid metabolism. Biochim Biophys Acta. 2013;1832(6):773–9.

52. Yennawar NH, Islam MM, Conway M, Wallin R, and Hutson SM. Human mitochondrial branched chain aminotransferase isozyme: structural role of the CXXC center in catalysis. J Biol Chem. 2006;281(51):39660–71.

53. He A, Chen X, Tan M, Chen Y, Lu D, Zhang X, et al. Acetyl-CoA Derived from Hepatic Peroxisomal β-Oxidation Inhibits Autophagy and Promotes Steatosis via mTORC1 Activation. Mol Cell. 2020;79(1):30–42.e4.

54. Wellen KE, Hatzivassiliou G, Sachdeva UM, Bui TV, Cross JR, and Thompson CB. ATP-citrate lyase links cellular metabolism to histone acetylation. Science. 2009;324(5930):1076–80.

55. Schug ZT, Peck B, Jones DT, Zhang Q, Grosskurth S, Alam IS, et al. Acetyl-CoA synthetase 2 promotes acetate utilization and maintains cancer cell growth under metabolic stress. Cancer Cell. 2015;27(1):57–71.

56. Han Y, Hu Z, Cui A, Liu Z, Ma F, Xue Y, et al. Post-translational regulation of lipogenesis via AMPK-dependent phosphorylation of insulin-induced gene. Nature Communications. 2019;10(1):623.

57. Jang C, Wada S, Yang S, Gosis B, Zeng X, Zhang Z, et al. The small intestine shields the liver from fructose-induced steatosis. Nat Metab. 2020;2(7):586–93.

58. Tian J, Goldstein JL, and Brown MS. Insulin induction of SREBP-1c in rodent liver requires LXRα-C/EBPβ complex. Proceedings of the National Academy of Sciences. 2016;113(29):8182–7.

59. Li S, Brown MS, and Goldstein JL. Bifurcation of insulin signaling pathway in rat liver: mTORC1 required for stimulation of lipogenesis, but not inhibition of gluconeogenesis. Proc Natl Acad Sci U S A. 2010;107(8):3441–6.

60. Horton JD, Goldstein JL, and Brown MS. SREBPs: activators of the complete program of cholesterol and fatty acid synthesis in the liver. J Clin Invest. 2002;109(9):1125–31.

61. Dentin R, Benhamed F, Hainault I, Fauveau V, Foufelle F, Dyck JR, et al. Liver-specific inhibition of ChREBP improves hepatic steatosis and insulin resistance in ob/ob mice. Diabetes. 2006;55(8):2159–70.

62. Erion DM, Popov V, Hsiao JJ, Vatner D, Mitchell K, Yonemitsu S, et al. The role of the carbohydrate response element-binding protein in male fructose-fed rats. Endocrinology. 2013;154(1):36–44.

63. Iizuka K, Bruick RK, Liang G, Horton JD, and Uyeda K. Deficiency of carbohydrate response element-binding protein (ChREBP) reduces lipogenesis as well as glycolysis. Proceedings of the National Academy of Sciences of the United States of America. 2004;101(19):7281–6.

64. Abdul-Wahed A, Guilmeau S, and Postic C. Sweet Sixteenth for ChREBP: Established Roles and Future Goals. Cell Metab. 2017;26(2):324–41.

65. Owen JL, Zhang Y, Bae S-H, Farooqi MS, Liang G, Hammer RE, et al. Insulin stimulation of SREBP-1c processing in transgenic rat hepatocytes requires p70 S6-kinase. Proceedings of the National Academy of Sciences. 2012;109(40):16184–9.

66. Yecies JL, Zhang HH, Menon S, Liu S, Yecies D, Lipovsky AI, et al. Akt stimulates hepatic SREBP1c and lipogenesis through parallel mTORC1-dependent and independent pathways. Cell Metab. 2011;14(1):21–32.

67. Peterson TR, Sengupta SS, Harris TE, Carmack AE, Kang SA, Balderas E, et al. mTOR complex 1 regulates lipin 1 localization to control the SREBP pathway. Cell. 2011;146(3):408–20.

68. Li Y, Xu S, Mihaylova MM, Zheng B, Hou X, Jiang B, et al. AMPK Phosphorylates and Inhibits SREBP Activity to Attenuate Hepatic Steatosis and Atherosclerosis in Diet-Induced Insulin-Resistant Mice. Cell Metabolism. 2011;13(4):376–88.

69. Wang Q, Jiang L, Wang J, Li S, Yu Y, You J, et al. Abrogation of hepatic ATP-citrate lyase protects against fatty liver and ameliorates hyperglycemia in leptin receptor-deficient mice. Hepatology. 2009;49(4):1166–75.

70. Ballantyne CM, Banach M, Mancini GBJ, Lepor NE, Hanselman JC, Zhao X, et al. Efficacy and safety of bempedoic acid added to ezetimibe in statin-intolerant patients with hypercholesterolemia: A randomized, placebo-controlled study. Atherosclerosis. 2018;277:195–203.

71. Ballantyne CM, McKenney JM, MacDougall DE, Margulies JR, Robinson PL, Hanselman JC, et al. Effect of ETC-1002 on Serum Low-Density Lipoprotein Cholesterol in Hypercholesterolemic Patients Receiving Statin Therapy. American Journal of Cardiology. 2016;117(12):1928–33.

72. Pinkosky SL, Newton RS, Day EA, Ford RJ, Lhotak S, Austin RC, et al. Liver-specific ATP-citrate lyase inhibition by bempedoic acid decreases LDL-C and attenuates atherosclerosis. Nature Communications. 2016;7(1):13457.

73. Ray KK, Bays HE, Catapano AL, Lalwani ND, Bloedon LT, Sterling LR, et al. Safety and Efficacy of Bempedoic Acid to Reduce LDL Cholesterol. New England Journal of Medicine. 2019;380(11):1022–32.

74. Ballantyne CM, Davidson MH, MacDougall DE, Bays HE, DiCarlo LA, Rosenberg NL, et al. Efficacy and Safety of a Novel Dual Modulator of Adenosine Triphosphate-Citrate Lyase and Adenosine Monophosphate-Activated Protein Kinase in Patients With Hypercholesterolemia. Journal of the American College of Cardiology. 2013;62(13):1154–62.

75. Uhlén M, Fagerberg L, Hallström BM, Lindskog C, Oksvold P, Mardinoglu A, et al. Tissue-based map of the human proteome. Science. 2015;347(6220):1260419.

76. Sjöstedt E, Zhong W, Fagerberg L, Karlsson M, Mitsios N, Adori C, et al. An atlas of the protein-coding genes in the human, pig, and mouse brain. Science. 2020;367(6482):eaay5947.

77. Karlsson M, Zhang C, Méar L, Zhong W, Digre A, Katona B, et al. A single cell type transcriptomics map of human tissues. Science Advances. 2021;7(31):eabh2169.

78. Xie J, Tai PWL, Brown A, Gong S, Zhu S, Wang Y, et al. Effective and Accurate Gene Silencing by a Recombinant AAV-Compatible MicroRNA Scaffold. Mol Ther. 2020;28(2):422–30.

79. Xie J, Ameres SL, Friedline R, Hung JH, Zhang Y, Xie Q, et al. Long-term, efficient inhibition of microRNA function in mice using rAAV vectors. Nat Methods. 2012;9(4):403–9.

80. Sena-Esteves M, and Gao G. Introducing Genes into Mammalian Cells: Viral Vectors. Cold Spring Harb Protoc. 2020;2020(8):095513.

81. Livak KJ, and Schmittgen TD. Methods. Academic Press Inc.; 2001:402–8.

82. Han X. Lipidomics. 2016:21–52.

83. Wang M, and Han X. Multidimensional mass spectrometry-based shotgun lipidomics. Methods Mol Biol. 2014;1198:203–20.

84. Han X, Yang K, and Gross RW. Microfluidics-based electrospray ionization enhances the intrasource separation of lipid classes and extends identification of individual molecular species through multi-dimensional mass spectrometry: development of an automated high-throughput platform for shotgun lipidomics. Rapid Commun Mass Spectrom. 2008;22(13):2115–24.

85. Yang K, Cheng H, Gross RW, and Han X. Automated Lipid Identification and Quantification by Multidimensional Mass Spectrometry-Based Shotgun Lipidomics. Analytical Chemistry. 2009;81(11):4356–68.

86. White PJ, Lapworth AL, An J, Wang L, McGarrah RW, Stevens RD, et al. Branched-chain amino acid restriction in Zucker-fatty rats improves muscle insulin sensitivity by enhancing efficiency of fatty acid oxidation and acylglycine export. Mol Metab. 2016;5(7):538–51.

87. Klett EL, Chen S, Edin ML, Li LO, Ilkayeva O, Zeldin DC, et al. Diminished acyl-CoA synthetase isoform 4 activity in INS 832/13 cells reduces cellular epoxyeicosatrienoic acid levels and results in impaired glucose-stimulated insulin secretion. J Biol Chem. 2013;288(30):21618–29.

88. Gao L, Chiou W, Tang H, Cheng X, Camp HS, and Burns DJ. Simultaneous quantification of malonyl-CoA and several other short-chain acyl-CoAs in animal tissues by ion-pairing reversed-phase HPLC/MS. J Chromatogr B Analyt Technol Biomed Life Sci. 2007;853(1-2):303–13.

89. Tomcik K, Ibarra RA, Sadhukhan S, Han Y, Tochtrop GP, and Zhang GF. Isotopomer enrichment assay for very short chain fatty acids and its metabolic applications. Anal Biochem. 2011;410(1):110–7.

90. He W, Wang Y, Xie EJ, Barry MA, and Zhang GF. Metabolic perturbations mediated by propionyl-CoA accumulation in organs of mouse model of propionic acidemia. Mol Genet Metab. 2021;134(3):257–66.

91. Dei Cas M, Paroni R, Saccardo A, Casagni E, Arnoldi S, Gambaro V, et al. A straightforward LC-MS/MS analysis to study serum profile of short and medium chain fatty acids. J Chromatogr B Analyt Technol Biomed Life Sci. 2020;1154:121982.

