## Supplemental Figures for "Paradoxical activation of SREBP1c and *de novo* lipogenesis by hepatocyte-selective ACLY depletion in obese mice"

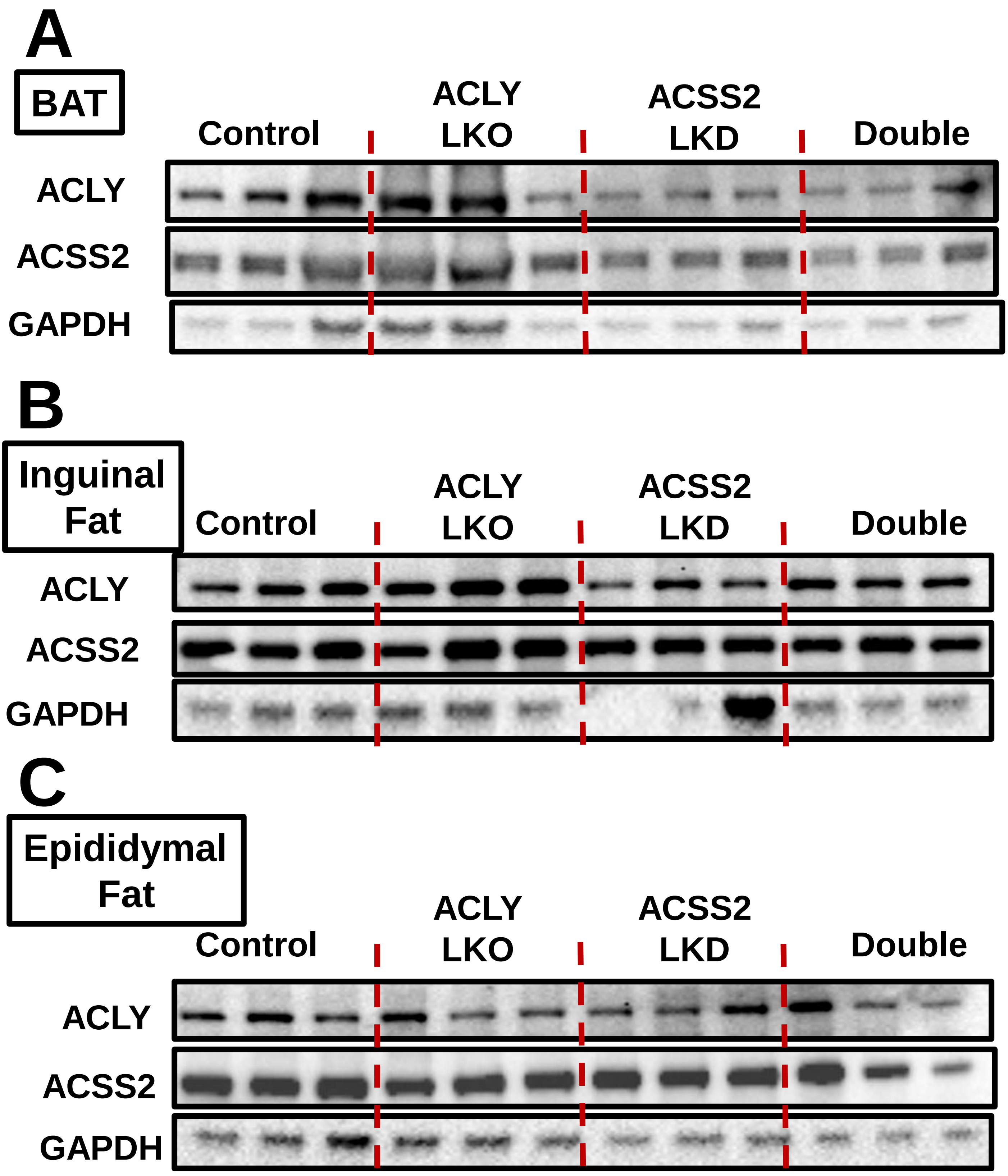


**Figure S1: Confirmation of liver specific targeting of the AAV constructs:** Immunoblots of ACLY and ACSS2 in **(A)** brown, **(B)** inguinal and **(C)** epididymal fat depots.

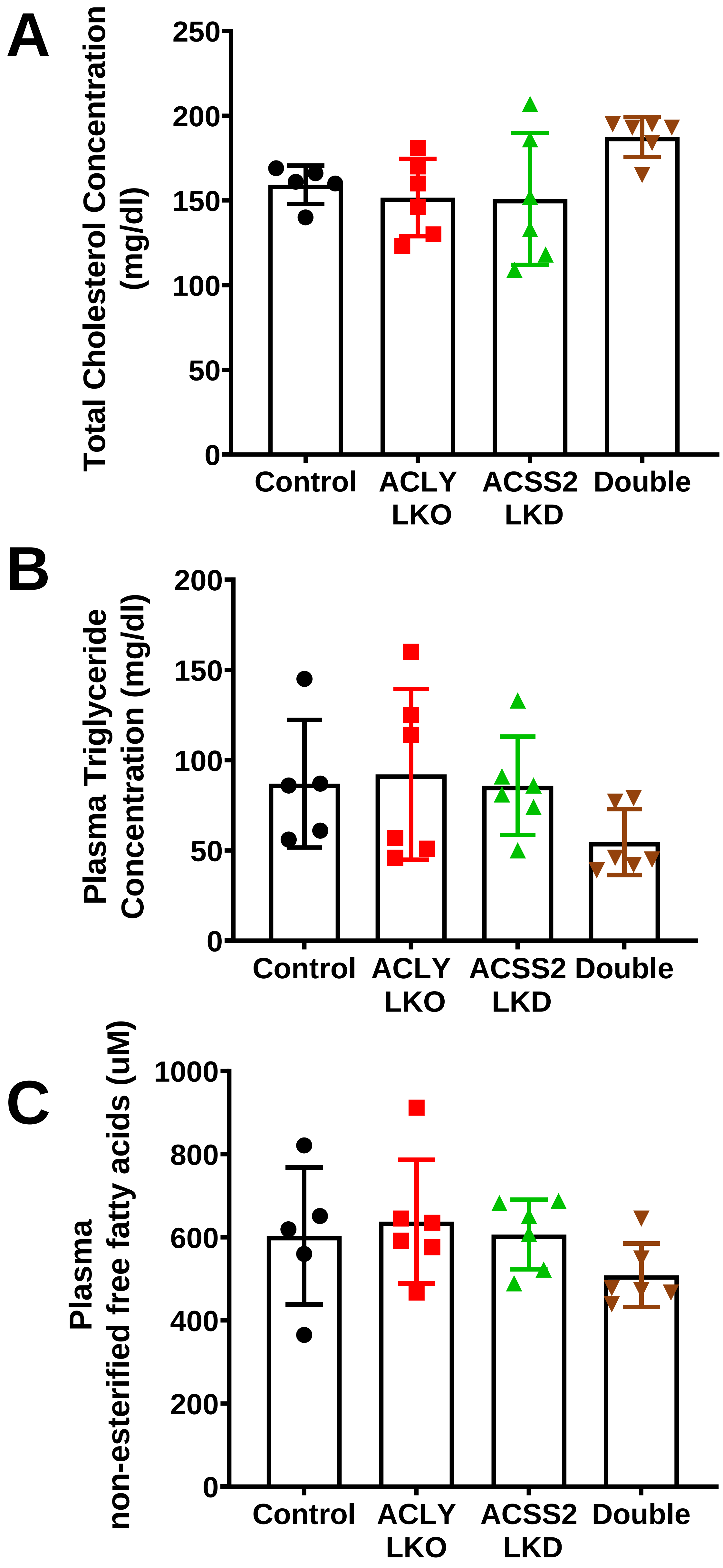


**Figure S2: Liver depletion of ACLY, ACSS2 or double depletion does not change the plasma lipid profile in HFD fed mice.** Plasma levels of **(A)** Total cholesterol, **(B)**TGs **(C)** Non esterified fatty acids.

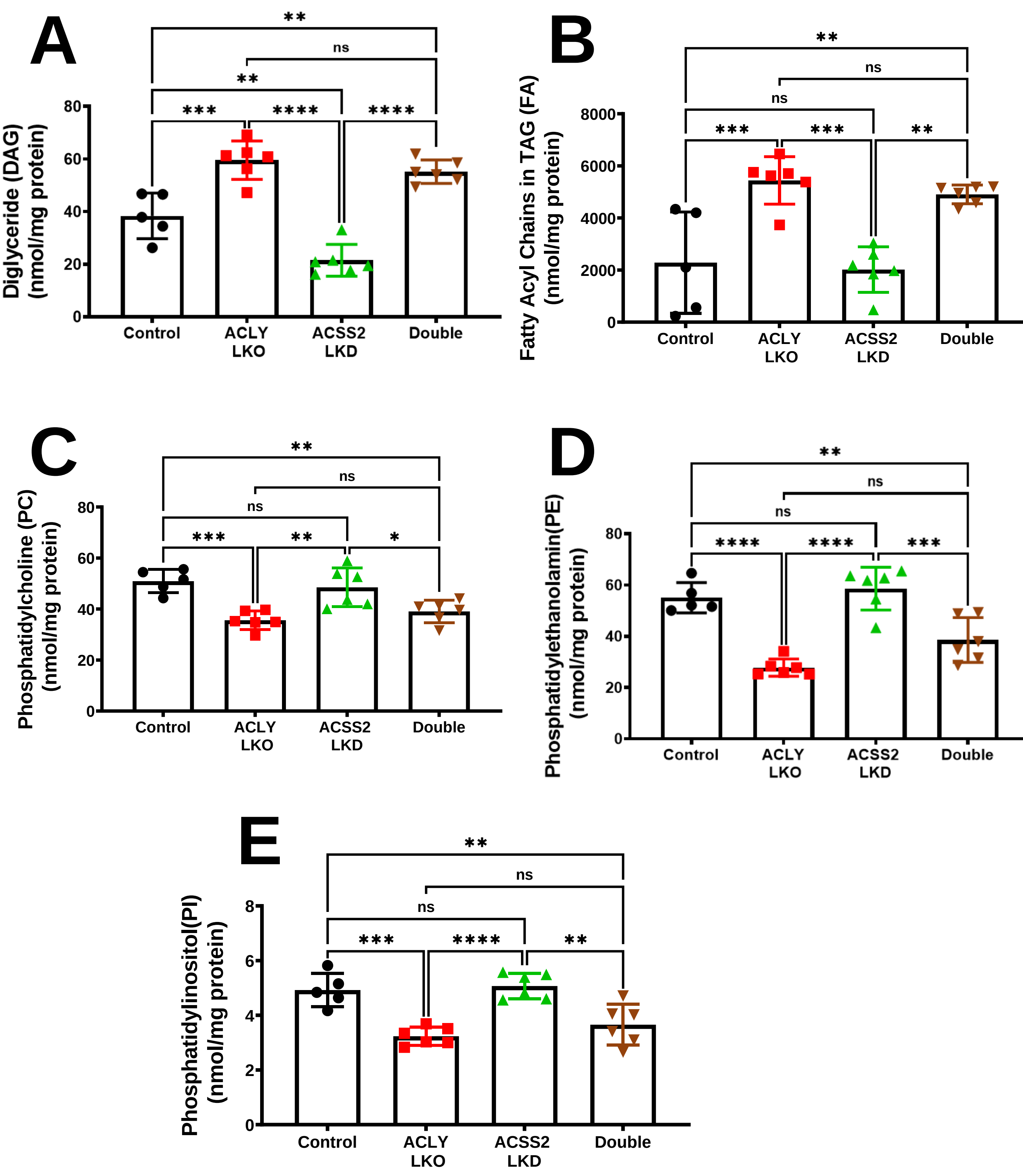


**Figure S3: Lipidomics analysis demonstrates accumulation of DAGs in ACLY-depleted but not ACSS2-depleted livers, and inverse correlation with phospholipid biosynthesis.** Liver **(A)** DAGs **(B)** Fatty acyl chains in TGs **(C)** Phosphatidylcholine **(D)** Phosphatidylethanolamine **(E)** Phosphatidylinositol levels in the livers of HFD fed groups. (ns: Not significant, *: $p<0.05$, **:$p<0.005$, ***:$p<0.0005$, ****:$p<0.00005$)
